# A Role for Astrocytic Insulin-Like Growth Factor I Receptors in the Response to Ischemic Insult

**DOI:** 10.1101/2023.01.13.523904

**Authors:** K. Suda, J. Pignatelli, L. Genis, A.M. Fernandez, E. Fernandez de Sevilla, I. Fernandez de la Cruz, A. Pozo-Rodrigalvarez, M. L. de Ceballos, S. Díaz-Pacheco, R. Herrero-Labrador, I. Torres Aleman

## Abstract

Increased neurotrophic support, including insulin-like growth factor I (IGF-I), is an important aspect of the adaptive response to ischemic insult. However, recent findings indicate that the IGF-I receptor (IGF-IR) in neurons plays a detrimental role in the response to stroke. Thus, we investigated the role of astrocytic IGF-IR on ischemic insults by deleting it using tamoxifen-regulated Cre deletion in glial fibrillary acidic protein (GFAP) astrocytes, a major cellular component in the response to injury. Ablation of IGF-IR in astrocytes (GFAP-IGF-IR KO mice) resulted in larger ischemic lesions, greater blood-brain-barrier disruption and more deteriorated sensorimotor coordination. RNAseq detetected increases in inflammatory, cell adhesion and angiogenic pathways, while the expression of various classical biomarkers of response to ischemic lesion, including aquaporin 4, complement 1q subunit a, early growth response protein 1, and C-C motif chemokine ligand 2, were significantly increased at the lesion site compared to control littermates. While serum IGF-I levels after injury were decreased in both control and GFAP-IR KO mice, brain IGF-I mRNA expression show larger increases in the latter. Further, greater damage was also accompanied by altered glial reactivity as reflected by changes in the morphology of GFAP astrocytes, and relative abundance of ionized calcium binding adaptor molecule 1 microglia. These results suggest a protective role for astrocytic IGF-IR in the response to ischemic injury.

## Introduction

Brain stroke is the second cause of death worldwide ^1^, and is a risk factor for dementia^2^. Unfortunately, current life-style, that increases its risks factors, dims hope for improvement ^3^. Albeit new therapeutic options are promising ^4, 5^, this condition is still a significant clinical burden demanding fresh insights if we want to reach better therapeutic outcomes ^6^. Endogenous neuroprotective pathways involving a myriad of homeostatic processes, that are highest during the subacute phase of response to ischemic damage ^6^, help restore function up to a certain degree after injury in many patients ^7^. Undoubtedly, a better knowledge of the processes involved in spontaneous recovery may lead to new therapeutic approaches by mimicking or potentiating them.

Insulin-like growth factor I (IGF-I) is considered a prototypical neurotrophic factor participating in protection against oxidative stress and inflammation ^8, 9^, two pathological cascades involved in stroke pathology ^10^. IGF-I has been reported beneficial in clinical studies of different brain illnesses ^11–14^, and has been directly linked to stroke outcome ^15, 16^. In this regard, different reports have shown that treatment of brain stroke with IGF-I is beneficial ^17–19^. Accordingly, IGF-I receptors (IGF-IR) are involved in IGF-I neuroprotection ^20^, including brain ischemia in infant mice ^21^. However, reduced levels of serum IGF-I ^22^ and serum GH/IGF-I ^23^, or inactivation of the neuronal IGF-IR ^24^, was shown to be also protective against ischemic injury in adult mice. Since timing is a key determinant in stroke protection^6^, and these studies included experimental manipulation prior to insult, the role of this neuroprotective pathway in the response to ischemia needs further clarification.

In the present study we have re-examined the role of IGF-IR in stroke injury following the same timing; i.e.: manipulating it prior to insult. We have found that in opposition to what was seen in mice with impaired IGF-IR activity in neurons ^24^, reducing IGF-IR activity in astrocytes resulted in increased damage after ischemic injury. Greater functional impairment and larger glial reactivity were linked with greater damage.

Based on these observations we suggest that the astrocytic IGF-IR is involved in neuroprotective responses to brain damage after ischemia and that this protection overcomes the apparent deleterious role of neuronal IGF-IR in the response to stroke.

## Methods

### Animals

Animals were kept under light/dark, (12 h/12h) conditions following EU guidelines (directive 86/609/EEC) and handled according to institutionally-approved procedures (Comunidad de Madrid, Proex 112/16). Animals were fed *ad libitum* with laboratory rodent chow and kept in standard laboratory cage conditions (4 mice/cage). All efforts were made to minimize suffering and to reduce the number of animals.

Transgenic mice used in this study have been characterized in detail before (Zegarra-Valdivia et al, in press). In brief, mice with tamoxifen-regulated deletion of IGF-IR in astrocytes (GFAP-IGF-IR KO mice) were obtained by crossing IGF-IR^f/f^ mice of B6, 129 background (IGF-IR^f/f^ mice; Jackson Labs; stock number: 012251) with CreERT2.GFAP mice of C57BL/6xSJL/J background (Jackson Labs, stock number: 012849). Mice lacking insulin receptors (IR) in astrocytes were obtained as described^25^, by crossing IR^f/f^ mice (B6.129S4(FVB)-Insr^tm1Khn^/J RRID:IMSR, Jackson labs; stock number 006955) with CreERT2.GFAP mice. Tamoxifen was injected for 5 consecutive days intraperitoneally (75 mg/kg, Sigma) at the age of 2 month, and animals were used a month later. GFAP-IGF-IR mice were treated with vehicle (corn oil). Using the tdTomato/eGFP reporter mouse to detect Cre-mediated deletion in response to tamoxifen administration in CreERT2. GFAP mice, we previously documented that it was restricted to astrocytes ^25^. Multiplex PCR for mouse genotyping included a common forward primer (P3, 5’–CTG TTT ACC ATG GCT GAG ATC TC–3’) and two reverse primers specific for the wild-type (P4, 5’–CCA AGG ATA TAA CAG ACA CCA TT-3’) and mutant (P2, 5’-CGC CTC CCC TAC CCG GTA GAA TTC–3’) alleles. GFAP-IGF-IR KO mice show reduced brain IGF-IR levels (Suppl Figure 1A), normal levels of serum IGF-I (Suppl Figure 1B) and normal brain size. This differs from mice lacking IGF-IR in neurons, that show high serum IGF-I levels ^24^. Further, levels of insulin receptor in GFAP-IGF-IR KO mice were normal (not shown). GFAP-IGF-IR KO mice were fertile, and food intake was also normal.

### Reagents

Anti-rabbit GFAP (1:1000) was obtained from DAKO (N0334), anti-chicken GFAP (1:1000) from ThermoFisher Scientific (PAI-10004), anti-rabbit Iba1 (1:1000) from WAKO (#019-19741), and anti-rabbit Aquaporin 4 (1:250) from Sigma (AB3594) were used. Anti-rabbit Alexa fluor 488, anti-rabbit Alexa fluor 647, and anti-chicken Alexa fluor 647 were used as secondary antibodies (ThermoFisher Scientific). Hoechst 33342 was purchased from Invitrogen (USA). 2,3,5-triphenyltetrazolium chloride (TTC) was purchased from Sigma (93140). Human IGF-I Quantikine ELISA Kit from R&D systems (DG100B) was used to measure IGF-I in serum, according to manufacturer’s protocol.

### Permanent brain ischemia

We followed previously reported methods of medial cerebral artery occlusion (MCAO), with modifications^26^. In brief, 6 to 9 months old mice (4-6 per group, both sexes) were anesthetized with 3% isoflurane (in O_2_) for induction and with 2% isoflurane for maintenance. Rectal temperature was maintained at 36.5 C with a heating pad. An incision perpendicular to the line connecting the lateral canthus of the left eye and the external auditory canal was made to expose and retract the temporalis muscle. A burr hole was drilled, and frontal and parietal branches of the MCA were exposed by cutting and retracting the dura. The frontal branch of the MCA was carefully disrupted. Following surgery, mice were returned to their cages, kept at room temperature and allowed free access to food and water. All physiological parameters measured: rectal temperature, mean arterial pressure and blood glucose levels were not different between groups. At various times after surgery, MRI, and SPECT/CT brain scans were performed, animals sacrificed and brains collected and processed for further analyses. To confirm the observations with MRI, 2,3,5-triphenyltetrazolium chloride (TTC) staining was done in an additional groups of mice, following procedures reported by others^27^. Sham surgery, that did not alter the brain parenchyma, was performed only in 2 control littermates to comply with reduced use of animals.

### Magnetic resonance imaging (MRI)

Brain damage was imaged by MRI 2 and 7 days after injury at 4.7 Teslas using a BIOSPEC BMT 47/40 (Bruker, Germany), equipped with a 12 cm actively shielded gradient system. Mice were anaesthetized and injected intraperitoneally with 0.4 mmol/kg Gadopentetate dimeglumine (Gadolinium, Magnevist, Germany). Mice were put in prone position inside a cradle to avoid unexpected movements. A respiration sensor was used to survey the animal’s vital functions. First, we acquired T2 weighted images using a fast spin echo sequence. The acquisition parameters were: TR = 4000 ms, effective TE = 60 ms, FOV = 3 cm, slice thickness = 1 mm and matrix = 256 × 192. This matrix size was increased during reconstruction by a zero-filling process in order to obtain images of 256 × 256 pixels. After that, weighted spin echo images were acquired (TR/TE=700/15 ms) using the same geometrical parameters as above. These images were used to calculate injury volume using ParaVision software (Bruker, Germany). All comparative studies between controls and experimental groups were performed at 1 week after injury since at this time we observed largest lesion sizes while at 2 days after injury brain edema interfered with lesion size measurement.

### Blood-brain-barrier (BBB) imaging

BBB integrity was analyzed using SPECT imaging with ^99^Technetium-diethylene-triamine-pentaacetate (Tc-DTPA; 900 μCi) 2 days after cerebral ischemia. Mice were iv injected with ^99^Tc-DTPA while awake and after 30 min of distribution of the tracer, they were imaged under anesthesia (2% isofluorane in O_2_), for 30 min (60 projections of 60 sec each), followed by CT acquisition (10 min), by means of an Albira II system (Bruker, Germany). SPECT and CT images were co-registered, reconstructed and analyzed. In normal conditions, the radiotracer is retained in the blood vessels, and it is only observed when the BBB is compromised. After fusion of both SPECT and CT images using the RM template Y.Uma, where the brain regions are outlined, images were masked for analysis of the volume presenting BBB disruption, to eliminate activity outside the brain (surgical damage). On those images, the activity to define the threshold was selected from the activity histogram. Then, the image was segmented by applying the “connected threshold”, localizing an initial “seed” where the maximal activity value is located, by the PMOD software v 3.3 (Bruker, Germany). The volume of interest (VOI) was outlined and copied to analyze the contralateral non-ischemic side of the brain, then both volumes (ischemic and that on the contralateral side) and the activity therein were assessed. The experimenter was blind to the mouse condition.

### Sensorimotor function

The adhesive removal test was performed 7 days after lesion as described ^28^. In brief, prior to lesion, we performed a protocol of adhesive removal training during 5 days, twice a day. Mice were familiarized to the testing room at least 30 min before starting the experiment to allow habituation to the new environment. Thereafter, animals were removed from their home cages and one adhesive tape strip was applied on each paw with equal pressure. Mice were placed in the testing box and animal’s behavior was observed. The time required to remove the tape strip from each paw was measured. Modified neurological severity scoring was performed 1 day after lesion, as described in detail elsewhere ^29^. Motor, sensory, and reflex responses were graded on a scale of 0 (normal) to 14 (maximal deficit).

### Cell cultures

Astrocyte cultures from P3 GFAP-IGF-IR KO and IGF-IR^f/f^ mice were prepared essentially as described ^30^. In brief, cells were grown in Dulbecco’s modified Eagle’s medium F12 (DMEM-F12) supplemented with 10% fetal calf serum. After 12 days, astrocytes were seeded at 2.5×10^5^ or 1.25×10^5^ cells/well in 6-well and 12-well culture plates, respectively. Astrocyte viability was determined by measuring cell metabolism with fluorescein diacetate (0.1 μg/ml FDA). FDA levels were measured in a plate reader (Fluostar, BMG, Germany) and expressed as arbitrary units. At least three independent experiments were done in 2-3 replicate wells.

### Immunocytochemistry

Immunostaining was performed as described ^31^. Animals were deeply anesthetized with pentobarbital and transcardially perfused with 4% paraformaldehyde in 0.1 M phosphate buffer, pH 7.4 (PB). Sagittal 50-μm thick sections were cut in a vibratome and collected in PB. Free-floating brain sections were blocked with 5% normal horse serum and incubated overnight at 4°C with the respective primary antibody in PB containing 0.1% bovine albumin, 3% horse serum, and 0.2% Triton X-100. After several washes in PB, sections were incubated with an Alexa-coupled secondary antibody (1:1000) and analyzed in a Leica microscope. Omission of primary antibody was used as control. Four mice were used for each group. Using a 20x objective, the ischemic border region of each animal was imaged with 2 slices, 1 field per slice. A total of 8 images were used for analysis.

### RNA sequence analysis

Cortical regions from ischemic GFAP-IGF-IR KO mice and littermates were dissected and total RNA isolated using Trizol Reagent and RNeasy mini kit (Qiagen). RNA was recovered in RNase free water (Ambion). Quantification and Quality assessment was performed using nanodrop (Thermofisher) and RNA integrity was evaluated by vertical electrophoresis (HS RNA cartridge, Bioptic SL). Only RNA samples with a RQ higher than 7 were used for posterior analysis. Libraries were prepared using TruSeq Total RNA Library prep (Illumina, Inc) and sequenced using NovaSeq600 (Illumina, Inc) following a Pair End (2×120bp) protocol. Only samples with a Coverage QC> 85% were considered as valid for analysis. NGS data were analyzed using R-based algorithms. First reads quality was assessed using FastQC tool. Sequence alignment was performed using Hisat2, and reads counting and differential expression analysis was performed using DESeq2. The Benjamini-Hochberg method of correction was established as the p-value adjustment method. Any gene with q-value >0.05 was considered as statistically significant, and as a differentially expressed gene (DEG). The R packages clusterProfiler and enrichplot were used to perform the Over Representation Analysis (ORA). The Gene Ontology (GO) knowledgebase was used as the background database, the Benjamini-Hochberg method of correction was set as the p-value adjustment method and the q-value cutoff at 0.05.

### Quantitative PCR

Total RNA isolation from cell lysates or brain tissue was carried out with Trizol. One μg of RNA was reverse transcribed using High Capacity cDNA Reverse Transcription Kit (Life Technologies) according to the manufacturer’s instructions. For the quantification of specific genes, total RNA was isolated and transcribed as above, and 62.5 ng of cDNA was amplified using TaqMan probes for vascular endothelial growth factor alpha (*Vegfa*) and *18S* as endogenous control (Life Technologies), and SYBR green probes for *cd93*, Platelet And Endothelial Cell Adhesion Molecule 1 (*pecam1*), Tenascin-C (*tnc*), *Igf1*, Early growth response protein 1 (*Egr1*), C-C Motif Chemokine Ligand 2 (*Ccl2*), nerve growth factor (*Ngf*), fibroblast growth factor 2 (*Fgf2*), brain-derived neurotrophic factor (*Bdnf*), aquaporin 4 (*Aqp4*), Complement C1q A Chain (*C1qa*) and glyceraldehyde-3-phosphate dehydrogenase (*Gapdh*) as endogenous control (Life Technologies). Each sample was run in triplicate in 20 μl of reaction volume using TaqMan Universal PCR Master Mix according to the manufacturer’s instructions (Life Technologies) or SYBR green Master Mix according to the manufacturer’s instructions (Life Technologies). All reactions were performed in a 7500 Real Time PCR system (Life Technologies). Quantitative real time PCR analysis was carried out as previously described ^32^. Results were expressed as relative expression ratios on the basis of group means for target transcripts versus reference reference transcript. All SYBR primers were designed by Sigma Aldrich (see Suppl. Table 1), and Taqman primers were designed by ThermoFisher Scientific. At least three independent experiments were done.

### Statistical analysis

Statistical analyses were carried out using GraphPad Prism 5 software or JMP 14 software. For in vitro assays, a minimum of 3 different experiments in duplicate/triplicate were done; the exact number of them are indicated for each case. The Kolmogorov-Smirnov test was used to check if the groups followed a normal distribution. If all groups passed the normality test, then we used one-way ANOVA followed by Bonferroni’s multiple comparison test. If the groups did not pass the normality test, we used a non-parametric Mann-Whitney test for comparing 2 groups, and the Kruskal-Wallis test (with the Dunn’s multiple comparison test as post-hoc) for more than two groups. Graphs depict mean value ± standard error (SEM) and the *p* values shown are coded as follows: *p<0.05, **p<0.01, ***p<0.001.

## Results

### Astrocytic IGF-I receptor-deficient mice are more vulnerable to ischemic damage

Since astrocytes participate in brain responses to ischemia ^33^ and astrocytic IGF-I has been shown to promote protection against stroke insult ^34^, we determined whether its receptor in this type of glial cell play a role in the response to ischemic damage. We used Cre/Lox mice where ablation of the IGF-IR in astrocytes was driven by tamoxifen (GFAP-IGF-IR KO mice). Ischemic injury in these mice was significantly larger one week after insult than control littermates, as determined by MRI analysis (Figure 1A, B). Increased lesion volume in GFAP-IGF-IR KO mice was histologically confirmed with TTC stainings (Figure 1C). The area under the curve of the unstained area, as determined by absence of TTC staining, was significantly larger than in littermates: GFAP-IGF-IR KO: 7.68 ± 1.08 mm^2^ vs control littermates: 2.68 ± 1.18 mm^2^, *p* < 0.05 (Figure 1D). The effect was specific for IGF-IR since elimination of the closely related insulin receptor (IR) in astrocytes using a similar tamoxifen-regulated GFAP-IR KO model ^25^ showed no obvious change in the ischemic area (Suppl Figure 2A, B).

**Figure 1:**
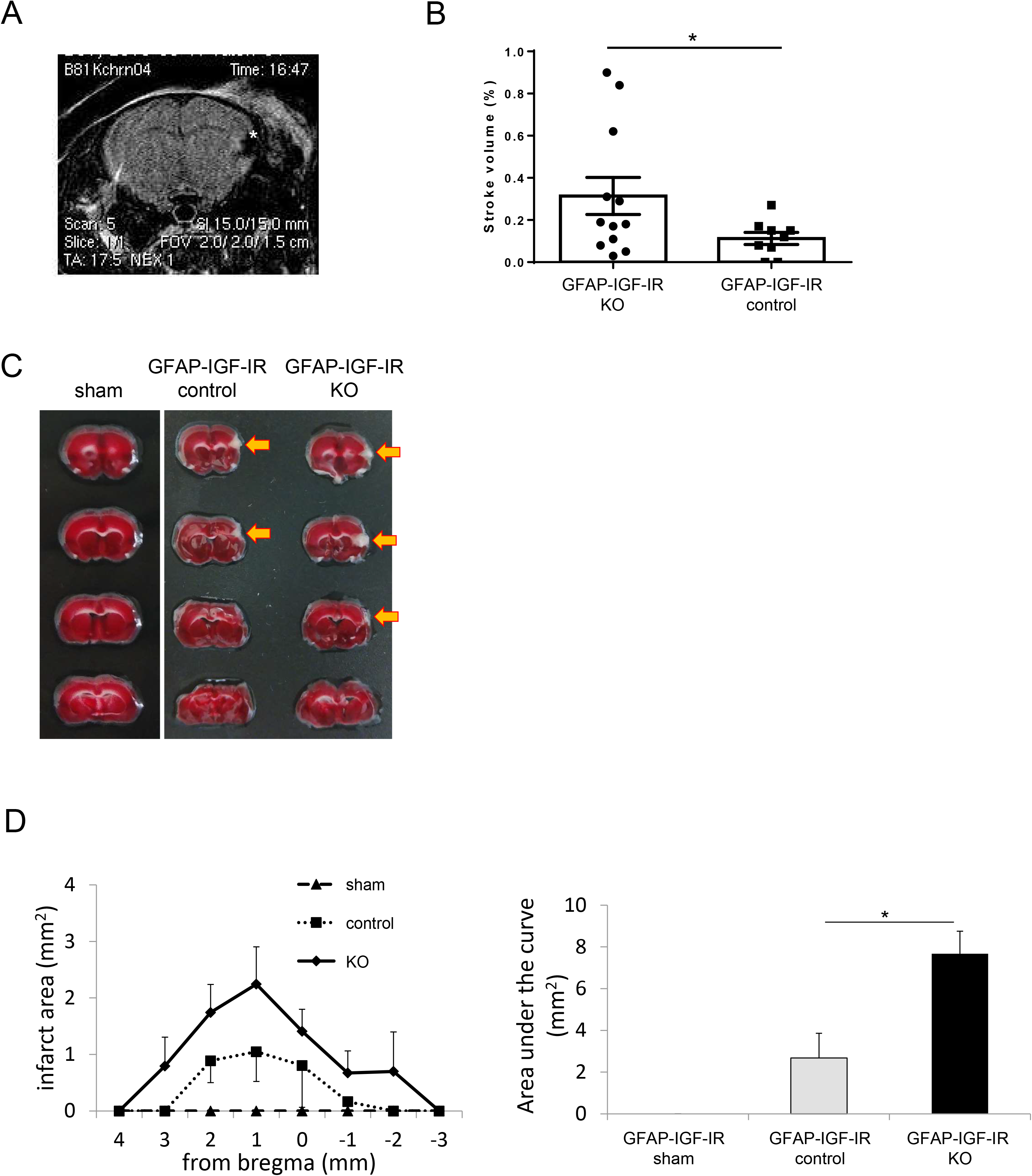
Reduced astrocytic IGF-IR activity and responses to ischemia. **A,** Representative brain magnetic resonance imaging (MRI); The asterisk shows the ischemic area. Note the absence of damage in the contralateral side. **B,** MRI evaluation of brain stroke volume in GFAP-IGF-IR KO mice (n=9) and GFAP-IGF-IR control littermates (n=12) (*p<0.05). **C,** Representative 2,3,5-triphenyltetrazolium chloride (TTC) staining in injured GFAP-IGF-IR KO and GFAP-IGF-IR littermates and in sham GFAP-IGF-IR littermates; arrowhead shows the ischemic area. **D,** Evaluation of stroke volume by TTC staining was performed in an independent group of mice using all brain sections throughout the anterior–posterior axis in GFAP-IGF-IR KO mice (n=5), GFAP-IGF-IR littermates (n=5), and GFAP-IGF-IR littermates after sham surgery (n=2, to reduce use of animals) confirmed greater damage in mutant mice. The evaluation of the area under the curve of these stroke volume is shown to the right (control vs. KO; *p*=0.047).

Injured GFAP-IGF-IR KO mice also show increased BBB disruption, as determined by ^99^DTPA SPECT/CT imaging (Figure 2A). Two days after ischemic insult, mean uptake of the tracer in the ipsilateral side of the brain was significantly higher in GFAP-IGF-IR KO mice, as compared to control littermates: GFAP-IGF-IR KO: 0.983 ± 0.067 vs control littermates: 0.764 ± 0.057 mm^2^, *p* < 0.05 (Figure 2B). However, at this early time, SPECT/CT analysis did not detect changes in the ischemic volume between experimental groups (Suppl Figure 2C).

**Figure 2:**
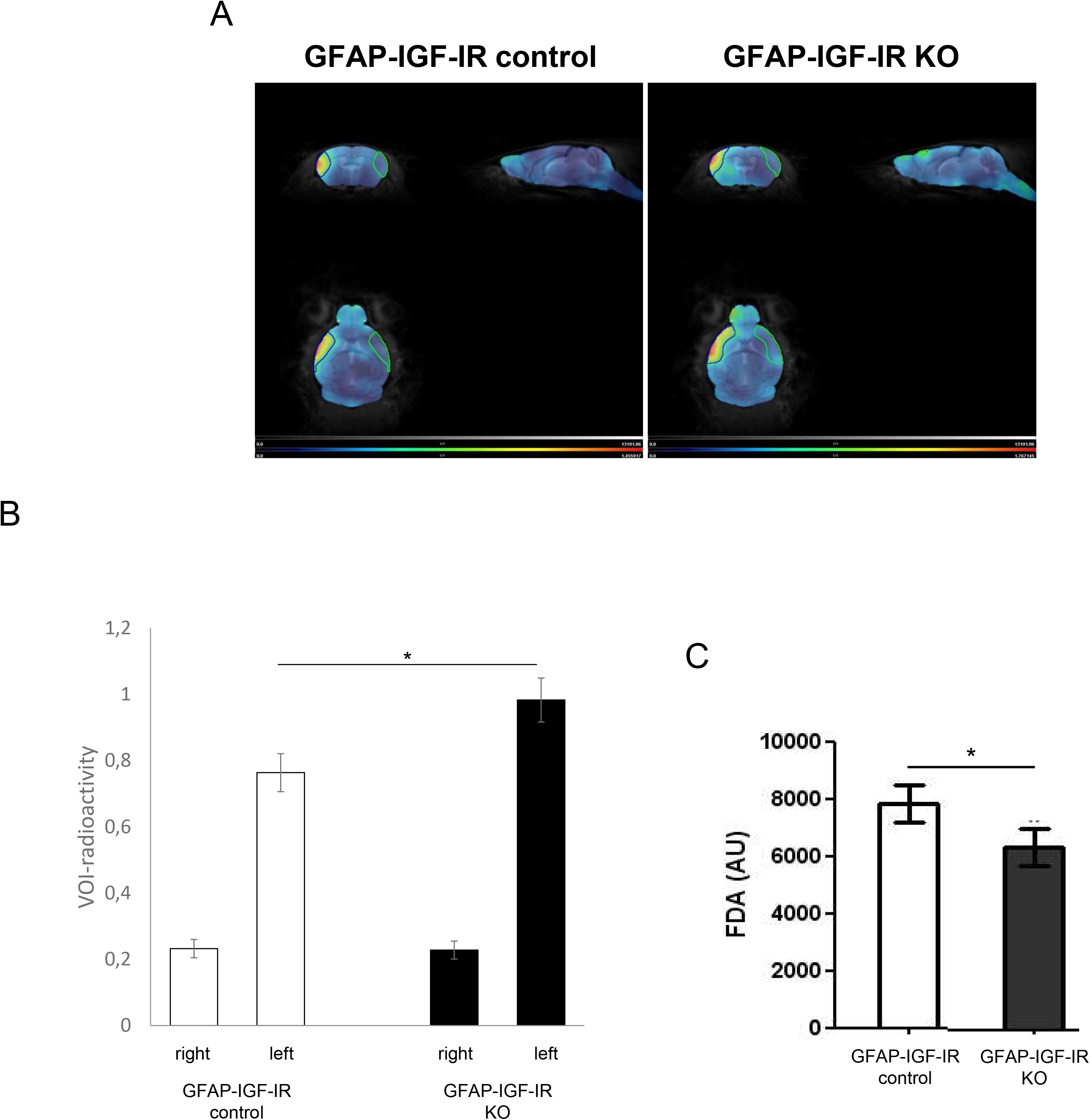
Reduced brain IGF-IR activity and BBB integrity. **A**, Representative SPECT/CT imaging with ^99^Technetium-diethylene-triamine-pentaacetate (Tc-DTPA); the outlined area is a volume of interest (VOI) and copied to analyze the contralateral non-ischemic side. **B**, Evaluation of the radioactivity by ^99^Tc-DTPA SPECT in GFAP-IGF-IR KO mice (n=5) and GFAP-IGF-IR control littermates (n=6) (control left vs. KO left; *p*=0.049). Radioactivity is expressed as % injected dose/cc. **C**, Evaluation of astrocyte viability determined by measuring cell metabolism with fluorescein diacetate (FDA); astrocytes were cultured from constitutively GFAP-IGF-IR KO mice or GFAP-IGF-IR control littermates (*p<0.05).

In accordance with these *in vivo* observations, astrocytes with ablated IGF-IR activity showed significantly reduced *in vitro* survival (Figure 2C), confirming previous observations that proper IGF-I signaling is required to support astrocyte survival ^30^.

### Functional impact of down-regulation of astrocytic IGF-IR activity in the response to brain ischemia

The functional impact of ischemic injury in GFAP-IGF-IR KO mice was assessed using two tests. Indeed, increased lesion site was reflected in greater neurological deficits; neurological severity scale in GFAP-IGF-IR KO was as follows: 6 mice scored 3, and 4 scored 2, while control littermates, 3 scored 2, 3 scored 1, and 1 scored 0 (*p* < 0.01; Figure 3A). As expected, injured control mice showed five times higher neurological severity score than littermates after sham surgery. Furthermore, one week after ischemic insult, mean time to remove the tape from the affected right paw in the adhesion removal test was longer in GFAP-IGF-IR KO mice than in control littermates: GFAP-IGF-IR KO: 191.8 ± 37.4 sec vs control littermates: 29.2 ± 10.9 sec, *p* < 0.05 (Figure 3B), reflecting poorer sensorimotor coordination. At this time after lesion, no changes in removal time were observed between control and littermates after sham surgery (Figure 3B), indicating that the functional impact in control injured animals is short-lasting.

**Figure 3:**
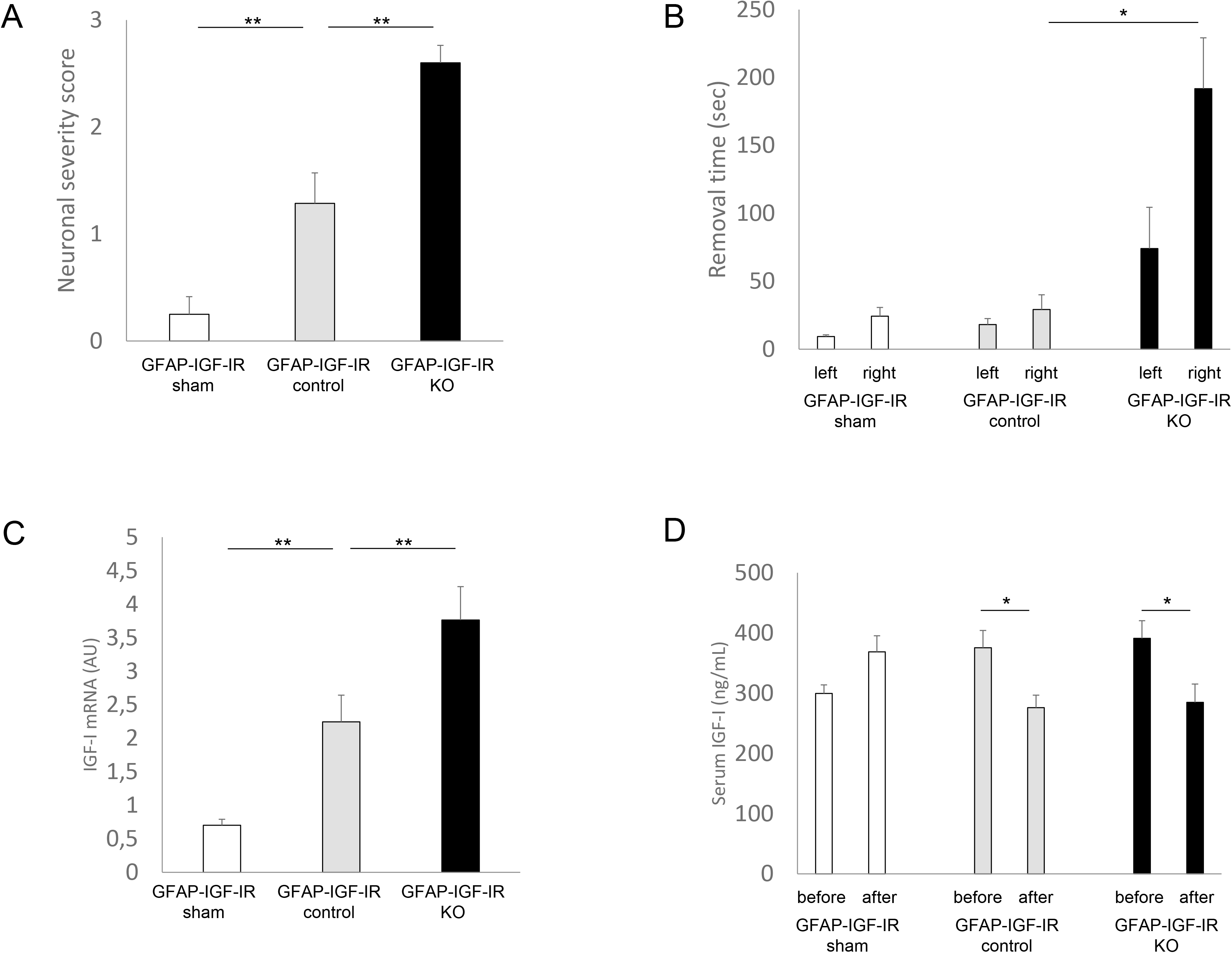
Functional impact of deletion of IGF-IR in astrocytes. **A,** Neurological severity score in GFAP-IGF-IR KO mice (n=10), GFAP-IGF-IR control littermates (n=7) (control vs. KO; *p*=0.006), and GFAP-IGF-IR control littermates after sham surgery (n=8) (sham vs. control; *p*=0.003). **B,** Ipsilateral sensorimotor function determined by the adhesive removal test in GFAP-IGF-IR KO mice (n=10), GFAP-IGF-IR control littermates (n=6), and GFAP-IGF-IR control littermates after sham surgery (n=4) (control right vs. KO right; *p*=0.022). **C,** IGF-I mRNA levels in ipsilateral cortex of GFAP-IGF-IR KO mice (n=6), GFAP-IGF-IR control littermates (n=6) (control vs. KO; *p*=0.010), and GFAP-IGF-IR control littermates after sham surgery (n=6) (sham vs. control; *p*=0.003). **D,** Serum IGF-I levels before and after injury; serum samples were collected from GFAP-IGF-IR KO (n=6) (KO before vs. after surgery; *p*=0.045), GFAP-IGF-IR control littermates before/after surgery (control before vs. after surgery; *p*=0.014), and GFAP-IGF-IR control littermates (n=6) before/after sham surgery.

After ischemic surgery, we observed increased gene expression of IGF-I in the ipsilateral cerebral cortex of injured control mice as compared to sham surgery mice, and even greater increased expression in injured GFAP-IGF-IR KO mice: GFAP-IGF-IR KO: 3.76 ± 0.50 vs control littermates: 2.25 ± 0.40 sec, *p* < 0.01 (Figure 3C). In parallel, serum IGF-I levels significantly decreased in both GFAP-IGF-IR KO and littermates after ischemic surgery (Figure 3D). This was paralleled by smaller weight gain in these groups (Suppl Figure 2D).

### Peri-lesion cellular and molecular adaptations to pre-injury reduced astrocyte IGF-IR

Under pathological conditions such as ischemia, astrocytes and other glial cells secrete inflammatory signals compromising BBB integrity ^35^. To determine the influence of the lack of IGF-IR in astrocytes prior to injury on adaptive responses, brains were collected for RNAseq gene expression analysis 3 days after surgery, when the expression of various inflammatory cytokines peaks in control animals^36–38^. The results show 22 known up- or down differentially regulated genes (DEGs) in GFAP-IGF-IR KO mice compared to controls with a q-value >0.05 and a fold change >0.5 Log2 (Figure 4A). Variability in expression levels of these 22 DEG show that control and GFAP-IGF-IR KO samples clustered depending only on genotype (Figure 4B). Gene ontology analysis (Figure 4C) indicated that the majority of DEG expressed in GFAP-IGF-IR KO mice were related to regulation of the inflammatory response (3 from 22 genes, fold enrichment 28), integrin cell surface integration (3 from 22 genes, fold change = 22.6), angiogenesis (4 from 22 genes, fold enrichment=12.8), and cell-cell adhesion (4 from 22 genes, fold change = 17.8). Pathway analysis showed those related to extracellular matrix interaction, cholesterol metabolism, and other pathways related to immune responses such as malaria, ALS, and leukocyte migration, and finally to the PI3-AKT signaling pathway (Figure 4D). Collectively, the results indicate that the immune response in GFAP-IGF-IR KO mice was increased after ischemic damage.

**Figure 4:**
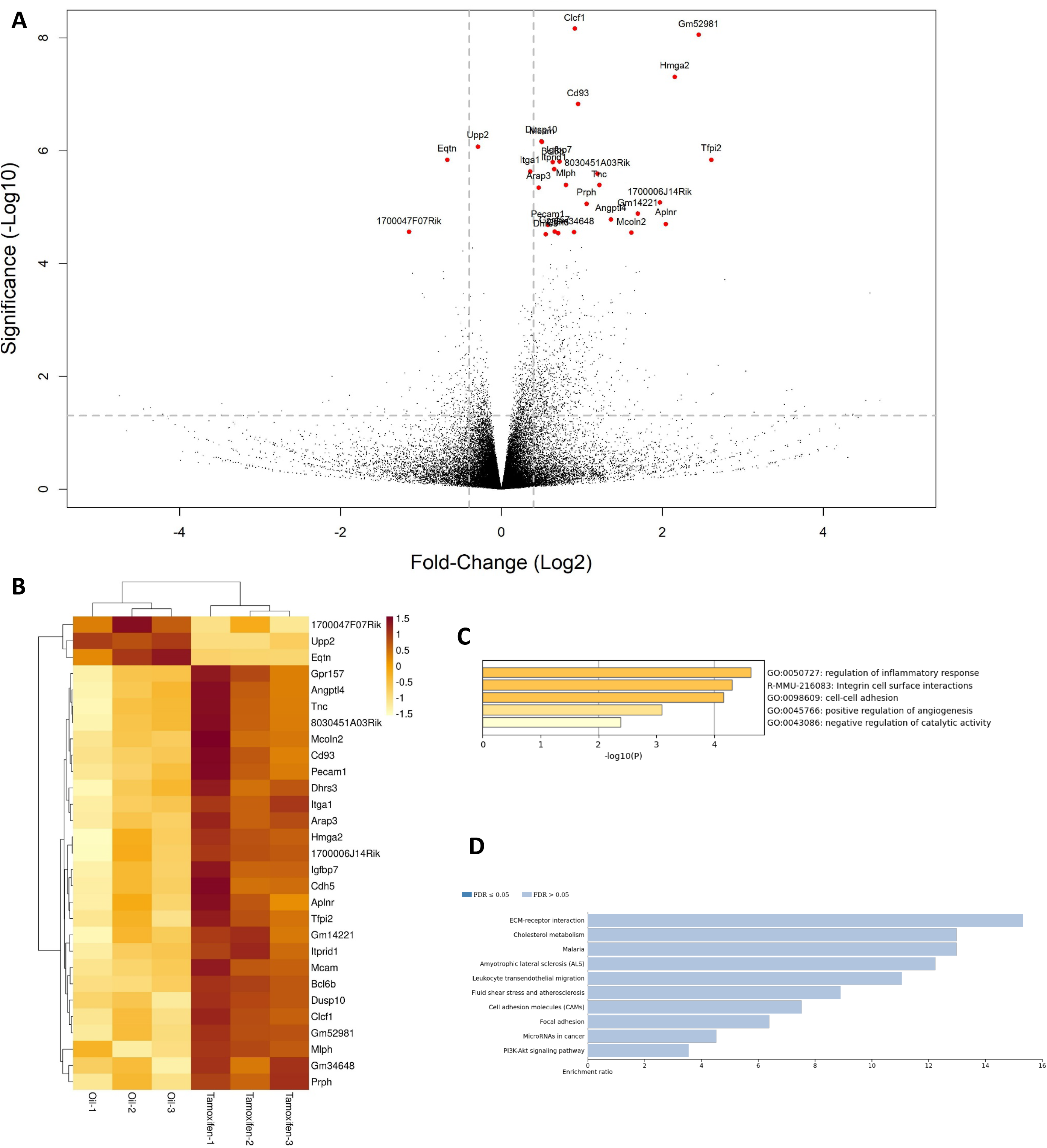
Transcriptional landscape after deletion of IGF-IR in astrocytes. **A,** Volcano plot showing differentially expressed genes between GFAP-IGF-IR KO mice and control littermates. **B,** Heatmap analysis of differentially expressed genes. **C,** Gene ontology analysis of differentially expressed genes. **D,** Gene ontology enrichment analysis of differentially expressed genes.

A few selected genes of these pathways: Cd93, Pecam1, and Tnc, were confirmed by qPCR (Figure 5A-C). We used the latter technique to examine expression of additional classical inflammatory factors known to be secreted by astrocytes and potentially affecting BBB integrity. Expression of Egr1, C1qa, Ccl2, and Aqp4 in the ipsilateral side of the cortex was significantly higher in GFAP-IGF-IR KO mice (Figure 5D-G) whereas the levels of various growth factors (Ngf, Fgf2, Bdnf, and Vegfα) were not significantly altered (Suppl Figure 3).

**Figure 5:**
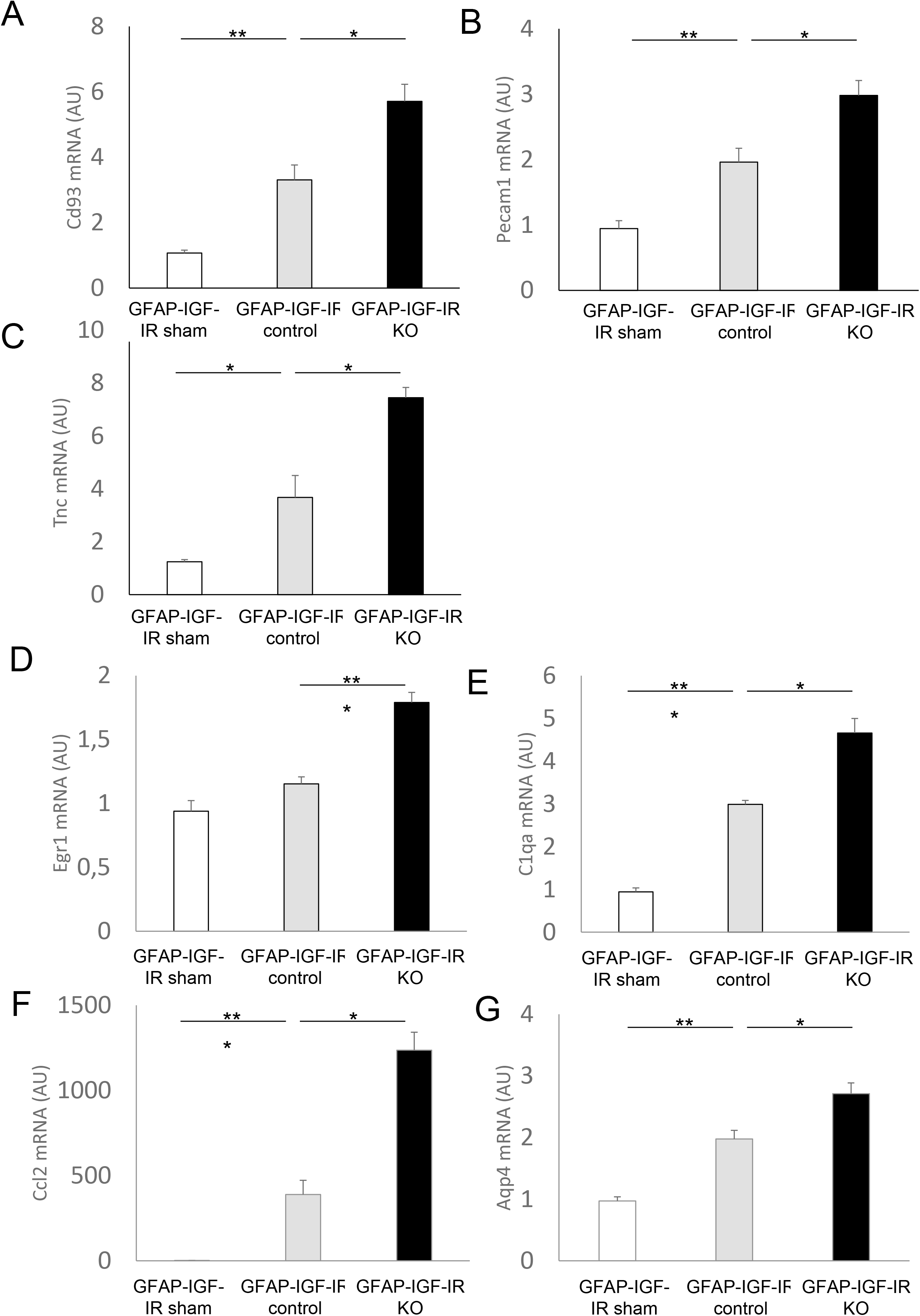
Molecular impact of IGF-IR deletion in astrocytes. **A,** mRNA levels of Cd93 (sham vs. control; *p*=0.001, control vs. KO; *p*=0.010). **B**, Pecam1 (sham vs. control; *p*=0.003, control vs. KO; *p*=0.013). **C**, Tnc (sham vs. control; *p*=0.013, control vs. KO; *p*=0.012). **D**, Egr1 (sham vs. control; *p*=0.207, control vs. KO; *p*=0.001). **E,** C1qa (sham vs. control; *p*=0.001, control vs. KO; *p*=0.020). **F,** Ccl2 (sham vs. control; *p*=0.001, control vs. KO; *p*=0.039). **G**, Aqp4 (sham vs. control; *p*=0.007, control vs. KO; *p*=0.038) in ipsilateral cortex of GFAP-IGF-IR KO mice (n=6), GFAP-IGF-IR control littermates (n=6), and GFAP-IGF-IR control littermates after sham surgery (n=6). *p<0.05, **p<0.01 and ***p<0.001 vs respective controls.

Next, we investigated whether worsened functional and anatomical outcomes were associated with changes in glial reactivity, a key cellular component in brain inflammatory responses. Immunostaining of glial fibrillary acidic protein (GFAP) astrocytes and allograft inflammatory factor 1 (Iba1) microglia showed that GFAP+ astrocytes neighboring the lesion area in GFAP-IGF-IR KO mice were as abundant as in control littermates in number, but show distinct morphological features (Figure 6A). Thus, the number of polarized GFAP+ astrocytes with cytoplasmic extensions towards the lesion site, that are considered to eventually form the peri-lesion scar^39^, was significantly higher in littermates: GFAP-IGF-IR KO: 227.67 ± 34.82 count/mm^2^ vs control littermates: 366.36 ± 24.11 count/mm^2^, *p* < 0.01 (Figure 6B), while the number of stellated astrocytes was significantly higher in GFAP-IGF-IR-KO mice: GFAP-IGF-IR KO: 398.64 ± 52.61 count/mm^2^ vs control littermates: 242.50 ± 41.22 count/mm^2^, *p* < 0.05 (Figure 6B). Furthermore, Iba1+ microglia positioned within 300 μm of the lesion area in GFAP-IGF-IR KO mice show an altered distribution (Figure 6C). Iba1+ cells in the proximal part of the lesion area, that is thought to be important in preventing spread of the injury ^40^, were comparatively reduced in GFAP-IGF-IR KO mice. Thus, microglia was widely distributed in the more distal part of the lesion in KO mice. As a result, the ratio of proximal/distal distribution of Iba1+ microglia was significantly reduced in GFAP-IGF-IR KO mice, compared to control littermates: GFAP-IGF-IR KO: 1.484 ± 0.082 vs control littermates: 2.422 ± 0.197, *p* < 0.001 (Figure 6D).

**Figure 6:**
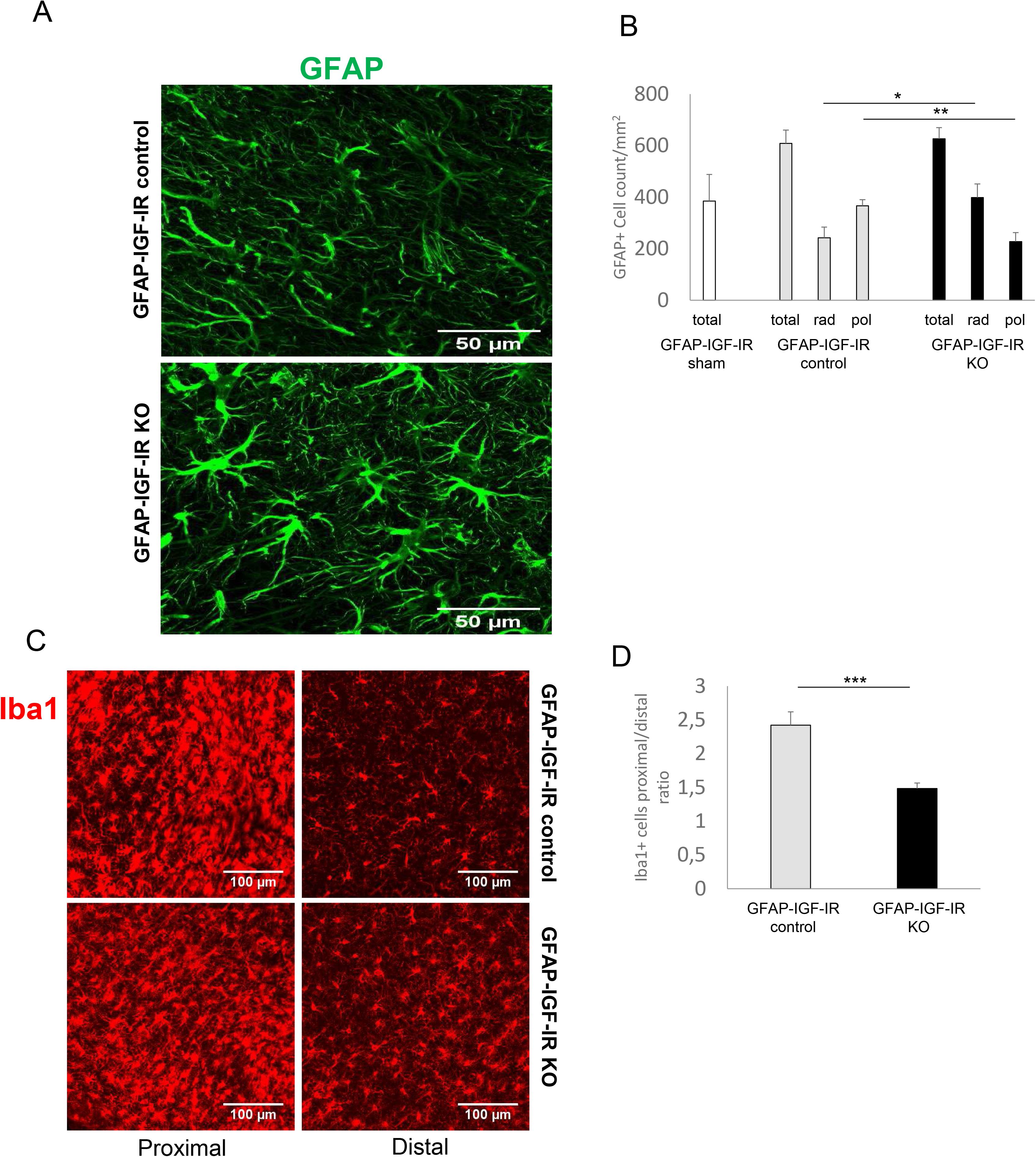
Differential morphological features in reactive astroglia and in microglia distribution in GFAP-IGF-IR KO mice after ischemic injury. **A,** Immunostaining of GFAP cells in GFAP-IGF-IR KO and GFAP-IGF-IR control littermates; the outlined area shows the representative polarized (upper) and radial cell (below). **B,** Evaluation of the number of GFAP+ cells with polarized elongated cytoplasm and radial morphology in GFAP-IGF-IR KO mice (n=8), GFAP-IGF-IR control littermates (n=8) (radial control vs. KO; *p*=0.046, polarized control vs. KO; *p*=0.008), and GFAP-IGF-IR control littermates after sham surgery (n=8). **C,** Immunostaining of Iba1+ cells in GFAP-IGF-IR KO and GFAP-IGF-IR control littermates; the left and right figures are taken within 300 μm (proximal) and beyond 300 μm (distal) from the lesion site, respectively. **D,** Evaluation of proximal / distal ratio of Iba1+ cell number in GFAP-IGF-IR KO mice (n=8) and GFAP-IGF-IR control littermates (n=8) (*p*=0.001).

In addition, we examined the cellular distribution of aquaporin (AQP) 4, which is present in astrocytic end-feet contacting cerebral vessels ^41^, since increased AQP expression is associated with exacerbation of cerebral edema during brain injury ^42^. We could not find a clear difference in the distribution of AQP4+ cells in GFAP-IGF-IR KO mice as compared to their controls (Figure 7A), although the AQP4+ area around the lesion site was significantly larger in GFAP-IGF-IR KO mice than in littermates: GFAP-IGF-IR KO: 0.185 ± 0.033 area/mm^2^ vs control littermates: 0.096 ± 0.017 area/mm^2^, *p* < 0.001 (Figure 7B).

**Figure 7:**
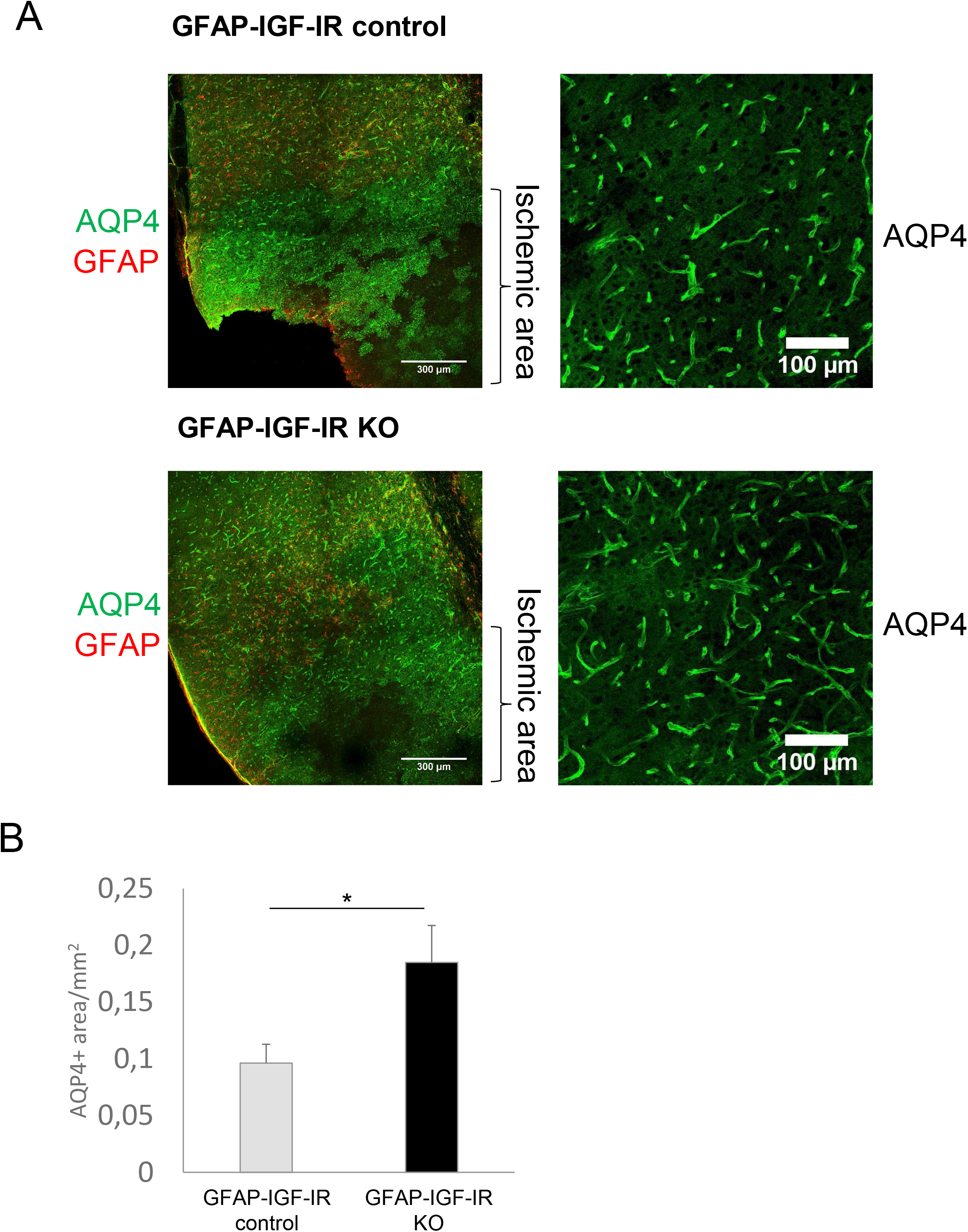
Differential features in reactive astroglia and aquaporin 4 expression in GFAP-IGF-IR KO mice after ischemic injury. **A**, Immunostaining of AQP4+ area in GFAP-IGF-IR KO and GFAP-IGF-IR control littermates; the right figures show high-resolution image taken around the lesion site. **B**, Evaluation of the AQP4+ area in GFAP-IGF-IR KO mice (n=8) and GFAP-IGF-IR control littermates (n=8) (*p*=0.036).

## Discussion

These results indicate that intact activity of the astrocytic IGF-IR is essential in reparative responses to ischemia as its absence produces greater tissue damage and more pronounced functional impairment. Thus, as already suggested for astrocytes in general ^43^, preservation of IGF-IR activity in this type of glial cells may prove a beneficial strategy for ischemia. Of note, reduced IGF-IR activity in neurons prior to injury was shown to reduce damage in a model of ischemia combined with hypoxia ^24^, suggesting either lesion-specific, cell autonomous actions of brain IGF-IR in the response to insult, or both. Diverse observations may help clarify these potentially conflicting results.

Neuronal IGF-IR deficient mice showed a phenotype that is quite different from GFAP-IGF-IR KO mice ^24^. The former show body overgrowth and high serum IGF-I levels, whereas the latter has these parameters within normal range. As indicated by the authors, neuronal IGF-IR deficient mice may show increased neuroprotection after injury through increased availability of IGF-I at the lesion site ^24^. Thus, besides the contribution of additional hypoxia exposure to permanent ischemic damage in their report^44^, defective IGF-IR signaling in astrocytes seems a key difference between the two types of mice. Indeed, when IGF-I signaling was specifically knocked down in astrocytes by using the tamoxifen-regulated strategy (similar to the one used to produce neuronal-specific mutants), lesion size after permanent stroke (similar to that used by De Magalhaes et al) was aggravated, while these mutant mice had normal body weight and unaltered serum IGF-I levels. Central and peripheral changes according to brain cell type where IGF-IR activity is modified, reinforce the notion of cell-specific actions of IGF-I.

The present observations suggest that cell autonomous actions of IGF-IR in brain cells may play varied roles in neuroprotection. Whereas opposite responses to ischemic damage between mice with reduced IGF-IR in neurons ^24^, and mice with reduced IGF-IR in astrocytes (present findings) could also be attributed to differences in the type of experimental model used in each study (plus/minus hypoxia), reducing IGF-IR activity in neurons protects not only against ischemia, but also against chronic conditions such as AD and spino-motor pathologies ^45–47^, making neuronal IGF-IR detrimental to responses to other types of tissue damage. Of note, since elimination of IGF-IR activity in astrocytes was sufficient to worsen responses to ischemia, a possible use of drugs blocking IGF-IR activity regardless of cell type for brain diseases ^48^, should be reconsidered. Moreover, the presence of IGF-IR in specific cellular compartments in glia ^49^ and neurons (in preparation), mediating complex neuronal plasticity patterns, at least in the latter ^50, 51^, suggests the existence of a highly compartmentalized role of this growth factor and its receptor on brain function. Indeed, neuroprotection by IGF-I against stroke differs when this growth factor is administered intra-cerebro-ventricularly ^17^, than when is overexpressed in astrocytes in the infarct area ^34^. As a corollary, the role of IGF-IR in other types of brain cells in the response to injury should be examined.

GFAP-IGF-IR KO mice showed cellular and molecular changes after ischemia compatible with aggravated responses to damage. Firstly, there was increased expression of a total of 20 genes involved in inflammatory, cell-adhesion and angiogenic pathways, which likely reflects potentiation of these putatively protective mechanisms when IGF-IR is absent in astrocytes. In particular, Cd93, which is involved in cell adhesion and clearance of apoptotic cells ^52^, Pecam1, a member of the immunoglobulin superfamily whose expression increases due to BBB disruption^53^, and Tnc, which is involved in neurogenesis^54^, are upregulated in GFAP-IGFIR-KO mice. However, abundant evidence points to complex patterns and timing of reparative responses after brain trauma already in normal mice, which combined with the absence of astrocytic IGF-I in our mice, makes it difficult to establish a direct link between these changes and the observed up-regulated genes. Additional limitations of our study are the lack of time-course analysis of molecular changes and the lack of comparison between male and female mice. Both limitations are due to the stringent ethical requirement of reducing the use of experimental animals. Future studies are now justified to provide time- and sex-dependent information in this model.

Reactive astrocytes in control mice showed elongated cytoplasmic extensions towards the lesion core 7 days after injury, while in GFAP-IGF-IR KO mice, astrocytes show a stellate morphology corresponding to an earlier, acute phase (1-4 days) of the process ^39^, suggesting a delayed/impaired response. Although the role of reactive glia after brain injury remains controversial ^55–58^, our observations indicate that morphological changes in reactive astrocytes lacking IGF-IR are associated to greater damage, suggesting that cellular interactions in the penumbra environment (probably directed to scar formation) are important modulators of post-injury responses ^59^.

Activated microglia in the ischemic brain is recruited to the penumbra region and is thought to prevent the spread of inflammation via phagocytosis of dead nerves ^40^, to exert neuroprotective effects in the acute phase of stroke^60^, while its processes are involved in BBB closure^61^. In GFAP-IGF-IR KO mice, microglia could potentially be more strongly recruited to the lesion site by higher local levels of Ccl2, which is predominantly secreted by astrocytes ^62^, but this was not the case; microglia in mutant mice was more abundantly present distal to the lesion site. We hypothesize that increased expression of C1qa, a microglial cytokine which contributes to phenotypic conversion of astrocytes into a neurotoxic subtype ^63^, may distort microglia recruiting. Our observation indicates that altered recruiting of microglia is associated to greater damage, suggesting that prevention of inflammation spread is impaired ^40^. Although further studies are needed to determine the mechanism by which activated microglia is not properly recruited to the vicinity of the lesion in the brain of GFAP-IGF-IR KO mice, an additional possibility might involve Egr1, which is strongly expressed in astrocytes during ischemia ^64^. Egr1 KO rats show decreased ischemic damage, edema improvement, and decreased number of recruited microglia, while the ischemic area is increased in rats overexpressing *Egr1* ^65^. In GFAP-IGF-IR KO mice, *Egr1* was highly expressed, so it can also contribute to altered distribution of microglia.

In GFAP-IGF-IR KO mice, expression of Aqp4 was increased, as determined by both qPCR and immunostaining. Aqp4 is a brain water channel affecting the evolution of cerebral edema after injury ^66^. While we did not observe a clear difference in the distribution of Aqp4, we speculate that an increased proportion of astrocytes with radial shapes found in GFAP-IR KO mice may interfere with proper distribution of Aqp4 in astrocytic end-feet and in this way hinder edema resolution. However, further analysis is needed.

Collectively, these findings support the use of targeted maintenance of IGF-IR activity in astrocytes as a therapeutic approach to improve resistance to ischemic injury. Indeed, these cells are important determinants of neurodegenerative processes ^67^, and key mediators of neuroprotection after ischemia ^68^. Therefore, strategies directed to this goal need to be developed. For example, physical exercise has been shown recently by us to increased total IGF-IR levels in mouse brain ^69^, and physical activity has been proven to protect against stroke in humans ^70^. It is possible that this protection involves the astrocytic IGF-IR.

## Acknowledgments

This work was funded by Ciberned (Instituto de Salud Carlos III, Spain) and CAM NEUROMETAB-CM (B2017/BMD-3700). We are thankful to M. Garcia and L Guinea for technical support and to the Omics Technologies Laboratory of the Cajal Institute.

## SUPPLEMENTARY FIGURE LEGENDS

**Supplementary Figure 1: A,** Brain IGF-IR levels in GFAP-IGF-IR KO mice and control littermates. **B,** Serum IGF-I levels in GFAP-IGF-IR KO (n=8) mice and littermates (n=6).

**Supplementary Figure 2: A,** Ischemic damage in GRFAP-IR KO mice assessed by TTC staining. **B,** Ischemic area in GFAP-IR KO mice was similar to control littermates, as assessed by TTC staining. **C,** Ischemic volume in GFAP-IGF-IR KO mice and controls estimated by ^99^DTPA SPECT/CT imaging. **D,** Growth curve of GFAP-IGF-IR KO male mice (n=5), GFAP-IGF-IR KO female mice (n=5), GFAP-IGF-IR male littermates (n=5), GFAP-IGF-IR female littermates (n=5).

**Supplementary Figure 3:** mRNA levels of Ngf (A), Fgf2 (B), Bdnf (C). and Vegfα (D) in injured GFAP-IGF-IR KO mice, control littermates with surgery, and sham littermates.

