## Supplementary figures and images for "A Role for Astrocytic Insulin-Like Growth Factor I Receptors in the Response to Ischemic Insult"

### Supplemental Figures

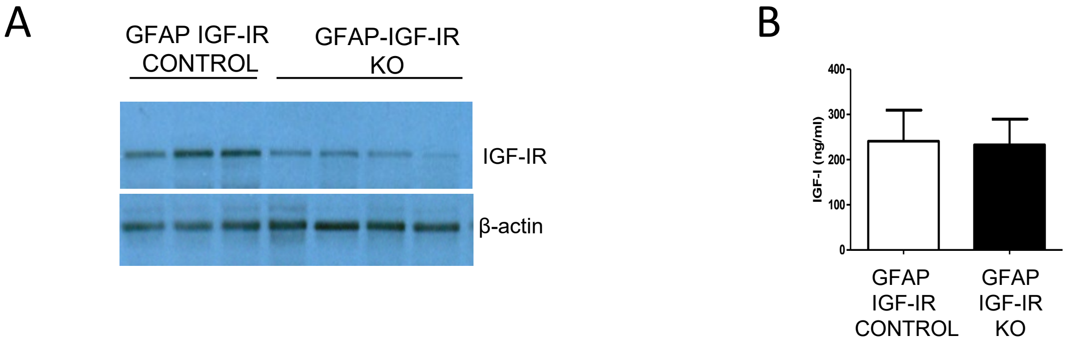


Supplementary Figure 1

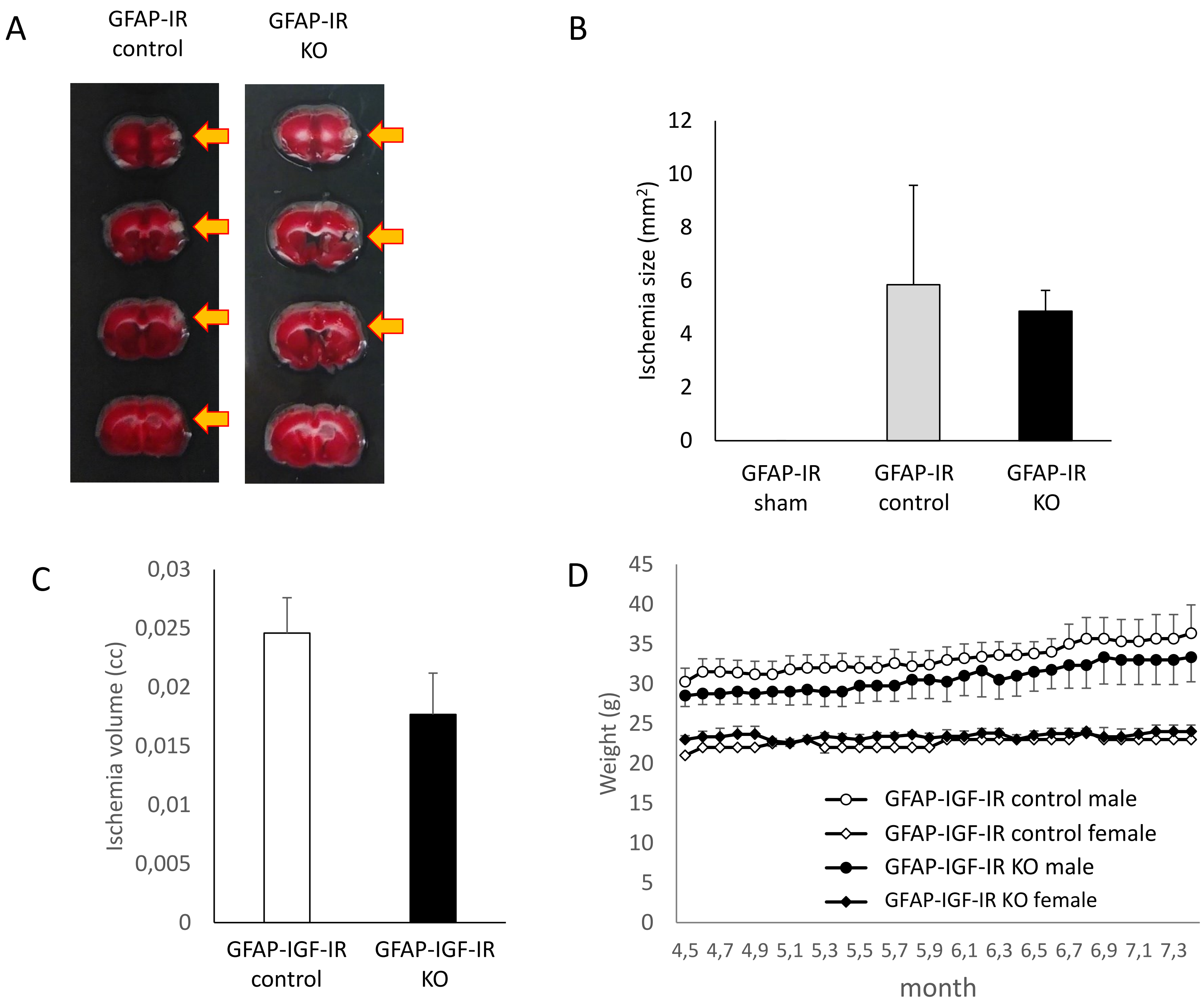


Supplementary Figure 2

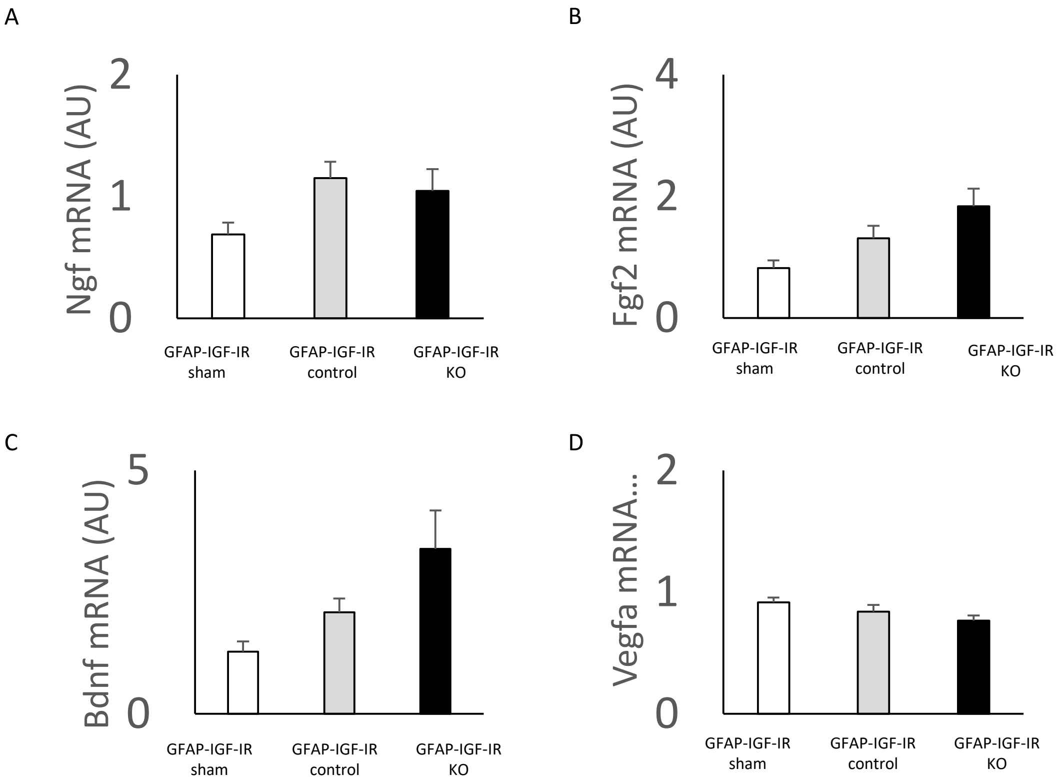


Supplementary Figure 3
